# Transcription can be sufficient, but is not necessary, to advance replication timing

**DOI:** 10.1101/2025.02.04.636516

**Authors:** Athanasios E. Vouzas, Takayo Sasaki, Juan Carlos Rivera-Mulia, Jesse L. Turner, Amber N. Brown, Karen E. Alexander, Laura Brueckner, Bas van Steensel, David M. Gilbert

## Abstract

DNA replication timing (RT) is correlated with transcription during cell fate changes but there are many exceptions and our understanding of this relationship suffers from a paucity of reductionist approaches. Here, we manipulated length and strength of transcription in hybrid-genome mouse embryonic stem cells (mESCs) at a single locus upstream of the silent, late replicating, Pleiotrophin (Ptn) gene, directly comparing RT to nascent transcription rates at engineered vs. wild-type alleles. First, we inserted four reporter genes that differ only in their promoter. Two promoters transcribed the reporter gene at high rates and advanced RT. The other two transcribed at lower rates and did not advance RT. Since these promoters may prove useful in applications where effects on RT are undesirable, we confirmed the inability of one of them to advance RT at numerous ectopic sites. We next juxtaposed these same four promoters upstream of the Ptn transcription start site where they all transcribed the 96kb Ptn gene and advanced RT to different extents correlated with transcription rates. Indeed, a doxycycline-responsive promoter, which could not advance RT when induced as a small reporter gene, elicited a rapid and reversible RT advance proportional to the rate of transcription, providing direct evidence that transcription itself can advance RT. However, deletion of the Ptn promoter and enhancer, followed by directed differentiation to neural precursors, eliminated induction of transcription throughout the entire Ptn replication domain, without preventing the switch to early replication. Our results provide a solid empirical base with which to re-evaluate many decades of literature, demonstrating that length and strength of transcription is sufficient but not necessary to advance RT. Our results also provide a robust system in which to rapidly effect an RT change, permitting mechanistic studies of the role of transcription in RT and the consequences of RT changes to epigenomic remodeling.

## INTRODUCTION

Mammalian chromosomes replicate segmentally in a temporal order, called the replication timing (RT) program. RT is cell-type specific and is disrupted in diseased states (Rivera-Mulia et al., 2017; Rivera-Mulia, Sasaki, et al., 2019), and the RT program is critical for the faithful propagation of a cell’s epigenomic state (Klein et al., 2021; Lande-Diner et al., 2009). In yeasts, RT is regulated by mechanisms that promote or antagonize steps in the conversion of DNA-bound MCM double hexamers into an active helicase (Hu & Stillman, 2023; Vouzas & Gilbert, 2023). Many of these mechanisms are conserved in mammalian cells, but the complexities of changing epigenomic states during cell fate transitions necessitate additional layers of upstream regulation. These additional layers regulate changes in the RT of sub-megabase replication domains (RDs) during cell fate transitions by altering the probability of initiating replication within multi-kilobase initiation zones (Dileep et al., 2019; Rivera-Mulia, Kim, et al., 2019; Sima et al., 2019; Vouzas & Gilbert, 2021; need to add Wang, Klein, Mol. Cell 2021). In mouse embryonic stem cells (mESCs), developmentally regulated early replicating RDs contain discrete early replication control elements (ERCEs) necessary to maintain domain-specific early RT, 3D architecture, chromatin compartment interactions, and transcription (Sima et al., 2019). These elements interact in 3D, are surrounded by acetylated histones and are bound by multiple transcription factors, which may function by creating a local sub-nuclear environment favorable for initiating replication (Sima et al., 2019; Turner et al., 2024). ERCEs are reminiscent of the Fkh1,2 transcription factors in budding yeast that interact and promote early replication independent of their transcription activity (Hoggard et al., 2021; Ostrow et al., 2017). RT can also be influenced by factors such as Rif1 that antagonizes initiation via mechanisms conserved with yeasts (Brueckner et al., 2020; Gnan et al., 2021; Heinz et al., 2018; Hiraga et al., 2017; Klein et al., 2021; Peace et al., 2014; Stow et al., 2022).

Transcriptional activation and silencing of genes is tightly coordinated with RT changes during cell fate transitions (Dileep et al., 2019; Hiratani et al., 2010; Rivera-Mulia et al., 2015) and early RT correlates with transcriptional activity in every multi-cellular organism investigated (Gilbert, 1986; Goldman et al., 1984; Hatton et al., 1988; Vouzas & Gilbert, 2021). However, the relationship between RT and transcription has been enigmatic. The correlation is largely driven by a majority of genes that are early replicating and expressed in all cell types (constitutively early housekeeping genes), while genes that are developmentally regulated show a much weaker correlation between RT and transcriptional activity (Rivera-Mulia et al., 2015). Indeed, some genes are induced when switching from early to late replicating (Hiratani et al., 2010; Rivera-Mulia et al., 2015). Moreover, mono-allelically expressed and asynchronously replicating genes show no relationship between their transcription and RT (Heskett et al., 2022).

Several groups have studied the effect of ectopic transcription on RT, with seemingly contradictory findings. In two studies targeting a strong acidic transcriptional activator to the promoter of late replicating genes in mESCs, both activated their transcription and advanced their RT (Brueckner et al., 2020; Therizols et al., 2014). In another case, targeting histone acetyltransferases to the mid S-phase replicating, human β-globin domain, inserted in the genome of transgenic mice, led to an advance in RT in the absence of transcriptional induction in both spleen lymphocytes and mouse embryonic fibroblasts isolated from these transgenic mice (Goren et al., 2008). Ectopic insertions of promoters have also had varied effects on RT. Insertion of an SV40-promoter driven vector in Chinese hamster cells was shown to advance RT of the insertion locus at one insertion site but not at two others (Gilbert & Cohen, 1990). Insertion of the β-actin and β-globin promoters in a mid-late replicating domain of chicken DT-40 cells advanced RT in a transcription dependent manner (Brossas et al., 2020). Also, in DT-40 cells, transcription of a short transcript or low levels of transcription of a longer transcript were unable to advance RT (Blin et. al., 2019; Hassan-Zadeh et. al.2012), while high levels of transcription of a long transcript did advance RT (Blin et. al.). The primary challenge with interpreting the seemingly contradictory results from the aforementioned collection of studies is the use of multiple methods to induce transcription and perturb RT in a variety of species, cell types and chromosomal sites.

Here, we take a systematic, reductionist, approach to dissect transcriptional activity in a late replicating domain with the express goal of identifying the elements of transcriptional activity that may have potential to advance RT of a developmentally regulated RD in which transcription of a single gene (Ptn) is induced coincident with a switch to early RT. We find that, with short selectable marker gene insertions, a high transcription rate can partially advance RT, although it also leads to a detectable number of longer transcripts that extend beyond the polyA termination site. However, lower levels of transcription, that are nonetheless fully capable of conferring drug resistance, do not affect RT in multiple vector and chromosomal contexts. Longer transcripts generated by the same promoters integrated at the same site but driving the 96kb Ptn gene, can all advance RT to varying extents. Taking advantage of the artificial Doxycyline-inducible promoter, we demonstrate that the advance in RT by this promoter, when driving Ptn, is proportional to the rate of transcription, is affected within hours of the onset of transcription and is reversible within a single cell cycle. However, we show that transcription of the Ptn gene is not necessary to advance RT in the context of differentiation; deletion of the enhancers and promoter of Ptn eliminated transcription induction without preventing the RT advance during a cell fate transition. Our results reconcile many seemingly contradictory reports in the literature, demonstrating that transcription can advance RT depending on the nature of the promoter and the length and strength of transcription but, is not necessary for an RT advance during a cell fate transition. Other factors must also be able to advance RT independent of transcription.

## RESULTS

### Transcription Per Se is Not Sufficient to Advance RT

We reasoned that one explanation for the varied effects of transcription on RT in the literature could be the use of different promoters. To systematically study the effect of transcriptional activation driven by different promoters on RT, we inserted a set of reporter genes into the same location upstream of the mouse Pleiotrophin (Ptn) gene, which encodes a neuronal growth factor. We identified the replication domain containing Ptn as one that switches from late to early coincident with transcriptional induction during the differentiation of mESCs to mouse neural precursor cells (mNPCs) (Hiratani et al., 2004, 2008, 2010). Subsequently, we and others have shown that the domain can advance RT in response to transcriptional induction induced by targeting a strong transactivator to the Ptn promoter in the absence of differentiation (Brueckner et al., 2020; Therizols et al., 2014). Ptn is the only gene in the domain that undergoes transcriptional induction at the time of the RT switch during differentiation to mNPCs; three other nearby genes, the Dgki gene whose 3’ end encroaches into the Ptn domain, an upstream non-coding RNA and the Chrm2 gene downstream of Ptn, all remain silent in both ESCs and mNPCs (Giorgetti et al., 2016; Rivera-Mulia et al., 2018). We constructed four reporter genes expressing the Hygromycin-Thymidine Kinase (HTK) gene fusion from a 2.0 kb transcript, differing only in the promoter used to drive expression (**Fig.1A**): the mouse and human phosphoglycerate kinase 1 (mPGK and hPGK) promoters, the cytomegalovirus immediate-early enhancer/chicken β-actin (CAG) promoter and the inducible tetracycline response element (TRE) promoter. The CAG promoter is a widely used synthetic construct that contains a cytomegalovirus (CMV) enhancer element, promoter, first exon and first intron of the chicken β-actin gene and the splice acceptor of the rabbit β-globin gene. The TRE promoter contains seven repeats of the bacterial tetO operator separated by a spacer. The mESC cell line used in all these studies expresses a reverse tetracycline transactivator (M2rtTA), consisting of the E. coli tetracyline repressor DNA binding domain fused to the Herpes virus VP16 acidic transactivation domain. The coding sequence for this artificial transactivator is integrated in front of and expressed constitutively from the Rosa26 promoter (Quinodoz et al., 2021). Upon introduction of Doxycycline (Dox), a derivative of Tetracycline, rtTA binds to the tetO elements of the TRE promoter and induces transcription. For all experiments, unless otherwise stated, cells with the inserted TRE promoter were treated with 2µg/ml of Dox for 24 hours. These reporter cassettes were inserted, one at a time, via CRISPR mediated homologous recombination, into a site ∼2.0 kb upstream of the Ptn gene transcription start site, avoiding all candidate cis-regulatory elements and replacing the genomic sequence marked as “YourSeq” in **Fig. 1B** (ENCODE Project Consortium et al., 2020). Insertion was made into the *musculus* allele of a *musculus* x *Castaneus* hybrid mESC line, greatly facilitating phasing of the two Ptn alleles due to the single nucleotide polymorphism (SNP) density of approximately one SNP every 150bp. In this way, we could distinguish the effects of insertion into one allele, while retaining the unmanipulated homologous allele in the same cells as an internal control, permitting the sensitivity to detect small RT changes (Rivera-Mulia et al., 2018; Sima et al., 2019).

**Figure 1:**
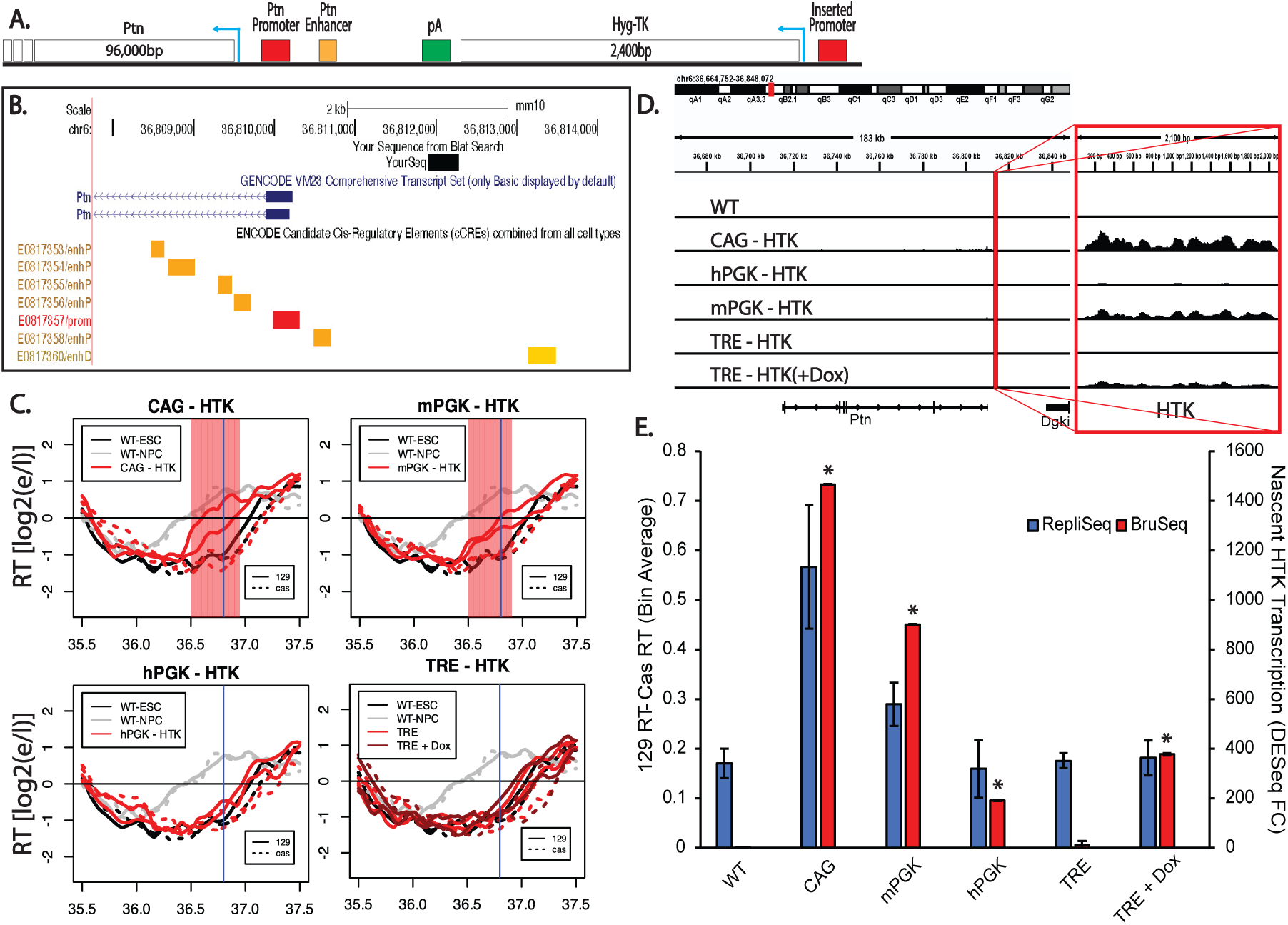
Promoter-specific effects of reporter gene insertion on RT. A) Schematic of the region directly upstream of the Ptn domain. All insertions were made ∼1.7kB upstream of the Ptn domain, leaving the natural Ptn promoter and enhancers intact. Arrows indicate the direction of the HTK and Ptn genes. The donor vectors used to make the insertions into the Ptn domain contained one of the four promoters used in this study driving the expression of a fusion positive/negative selectable marker consisting of the hygromycin resistance gene fused to the tyrosine kinase gene (HTK), followed by a transcription termination sequence (pA). Therefore, the inserted sequences were identical except for the promoter, allowing us to directly compare the effects of the four promoters on RT. B) UCSD Genome Browser track indicating the locations of the exact genomic sequence (YourSeq) that was replaced by the insertions, as well as the Ptn promoter (red), proximal enhancers (orange), and distal enhancer (yellow) elements. All candidate cis-regulatory elements remained intact after transgene replacement. C) RT of the Ptn domain for the CAG-HTK (top left), mPGK-HTK (top right), TRE-HTK (bottom left) and hPGK-HTK (bottom right) cell lines. In each plot, the black and gray lines indicate the RT of WT ESCs and mNPCs respectively. The red lines indicate the RT of the insertion cell line. In the TRE-Ptn plot, dark red lines and light red lines indicate the RT of TRE-Ptn cells with and without Dox, respectively. Solid and dashed lines indicate the RT of the *musculus* (mouse strain 129) and *castaneus* alleles, respectively. Vertical blue line indicates the site of the Promoter-HTK insertion. The red shading indicates 50kb windows with statistically significant differences in RT between WT casteneus and modified 129 alleles. D) Genome browser view of the Ptn domain displaying nascent RNA (Bru-Seq) signal for each one of the four promoters. The left track displays the unparsed Bru-Seq signal in the Ptn domain. The vertical red line indicates the locus where the HTK insertion was made. The zoomed in Bru-Seq signal for HTK is displayed on the right track. The dynamic range for both tracks is the same. Replicate experiments are shown in Supplemental Figure 1. E) Bar graph of the relationship between nascent HTK transcription and the observed advance in the RT of the Ptn domain. The asterisks indicate a significant difference in the levels of HTK expression calculated using DEseq, relative to HTK expression in the parental cell line, which is expected to be zero, since the parental line does not have an HTK insertion. The RT advance of the Ptn domain is measured as the average difference between the RT of 50kb bins of the *musculus* allele and the *castaneus* allele. As indicated, the *musculus* (129) allele replicates slightly earlier than the *castaneus* allele in WT; RT differences between the two alleles are normalized for these allelic differences. The error bars show the range of the difference among the 50kb bins between the two replicate experiments.

To assess the effects of these insertions on RT and transcription rates, we performed Repli-Seq and Bru-Seq (Marchal et al., 2018; Paulsen et al., 2014). The reporter cassettes harboring the CAG and mPGK promoters led to advances in the RT of the Ptn domain (**Fig. 1C**). In contrast, the reporter cassettes with hPGK or the inducible TRE promoter, with or without 2µg/ml Doxycycline (Dox) for 24 hours, were unable to advance the RT of the Ptn domain (**Fig. 1C**). Since all reporter cassettes transcribe HTK and sustain Hygromycin resistance, transcription of a reporter gene per sé is clearly not sufficient to advance RT. However, the CAG and mPGK promoters that were able to advance RT induced higher expression levels of HTK than the hPGK and TRE promoters (**Fig. 1D**). Note that Bru-Seq detects a small amount of transcriptional readthrough of the small transgene with the CAG and mPGK promoters, suggesting that the poly-A termination site, used in many vector designs, is not fully effective at constraining high levels of transcription to the vector sequences (**Fig. 1D, Sup. Fig. 1A-B**), albeit the level of read through Ptn expression is substantially lower than the levels of HTK expression **(Sup. Fig. 1C)**. Directly comparing the RT advance of the Ptn domain and the levels of transcription of HTK (**Fig. 1E**) revealed that, for the two promoters that did advance RT, the RT advance was greater when transcription rate was higher. One possible interpretation of these data is that there may be a transcription level threshold necessary to advance RT. Alternatively, the readthrough transcription with CAG and mPGK, creating a longer transcript, may elicit the RT advance **(Sup. Fig. 1D)**. Another possibility is that there are other, transcription-independent, differences between these promoters that influence their ability to advance RT. In summary, our results reconcile previous contradictory reports in the literature (Blin et al., 2019; Brossas et al., 2020; Gilbert & Cohen, 1990; Goren et al., 2008; Therizols et al., 2014)by demonstrating that some selectable marker gene insertions can advance RT while others cannot, and this is dependent upon the choice of promoter driving transcription, possibly due to transcription rate.

### The inability of hPGK to advance RT is vector and position independent

The hPGK promoter is commonly used to drive transcription and clearly produces sufficient gene expression for drug resistance, yet neither of these promoters can advance RT at the Ptn locus. If its lack of effect on RT is position-independent, such a promoter can be a useful tool to manipulate genomes without the concern of altering RT and consequently inducing significant changes in surrounding chromatin context. To determine whether hPGK can drive gene expression without affecting RT in a vector and position-independent manner, we analyzed the RT of hPGK promoter insertions in two previously characterized F121-9 clonal cell lines (Brueckner et al., 2020) with a total of 107 single-copy/single-allele insertions of a PiggyBac (PB) vector expressing GFP from the hPGK promoter (**Fig. 2A**). Although there was a slight bias towards PB integration into early replicating domains, we detected integrations at sites replicating throughout S phase, including both constitutive and developmental late replicating regions (**Fig. 2B**). Results revealed that all single-copy/single-allele insertion sites showed a high correlation between the *musculus* and *castaneus* alleles both in the mutant CM lines (**Fig. 2C**) and the parental WT cell lines **(Sup. Fig. 2A)**, demonstrating preservation of RT despite ectopic transcription in one allele. The few domains whose RT differed between alleles were at sites where there was a natural difference in the RT of the *musculus* vs *castaneous* genomes (**Fig. 2D**). Additionally, there were no statistically significant advances in the RT of all but of the insertion sites **(red line in Sup. Fig. 2B)**. The RT plot of the insertion site with the statistically significant RT advance reveals a modest RT advance (**Sup. Fig. 2C** – Top Right). A small number of other insertion sites had small but statistically non-significant RT changes **(Sup. Fig. 2C)**.

**Figure 2:**
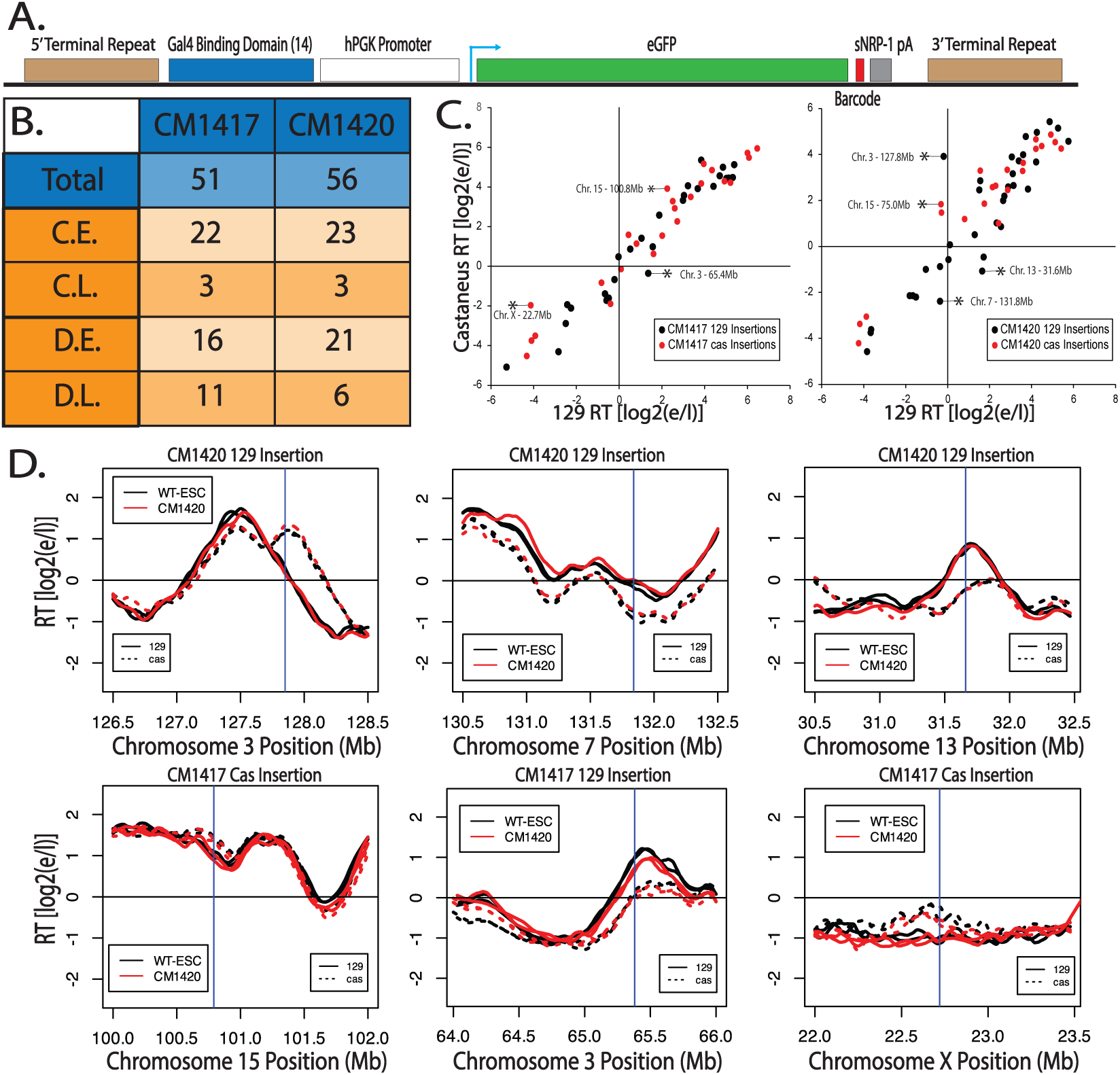
The absence of RT advance with hPGK vectors is not vector or position dependent. A) Schematic of the donor construct used for the Piggy Bac (PB) transposable element insertions, described in Brueckner et. al. (2020). The human PGK promoter is driving the expression of the enhanced Green Fluorescent Protein (eGFP) gene, along with a barcode specific to each insert. Transfection of this vector into F121-9 hybrid mESCs, along with the PB transposase, results in its insertion into multiple genomic sites. B) Total number and RT distribution of PB insertion sites in two clones CM1417 and CM1420. Piggy Bac insertions are slightly enriched in constitutively (C.E.) and developmentally (D.E.) early replication domains, vs. constitutively (C.L.) and developmentally (D.L.) late domains. C) Scatter plots of the RT of all ectopic insertion sites of the *musculus* (129) allele vs. *castaneus* allele for the CM1417 (left) and the CM1420 (right) clones. Black and red dots represent loci where the insertion was made in the 129 and cas allele respectively. Asterisks indicate insertion sites where there is a significant difference between the replication timing of the 129 and the cas alleles. Comparison of insertions to WT RT on the same alleles is shown in Supplemental Figure 2. D) RT profiles of six insertion sites, marked with asterisks in Fig. 2C, in which the two alleles replicate asynchronously, that replicate at different times demonstrates that their differences in RT are species specific, not due to the PB insertion. Vertical blue lines indicate the site of the PiggyBac insertion.

To assess the transcription levels at each insertion site we mined the mRNA-seq data from Brueckner et. al.. Since the inserted eGFP sequence is identical at all insertion sites this was challenging. However, dividing the total eGFP expression by the number of insertions in each clone reveals that the average eGFP expression is higher than 99% of all genes in both clones **(Sup. Fig. 2D)**. To assess expression levels across each insertion, we filtered for reads containing the entire 16 nucleotide barcode. It is important to note that coverage across any given 16 nucleotide segment of the genome may be low and therefore, this type of analysis can only be informative to assess the expression of barcodes relative to each other and not to other genes. Further, low or no expression of a barcode may be the result of low coverage, rather than a biological observation. Barcode specific expression analysis was limited to 61/107 insertions for which the barcode has been identified. Calculating the RPM of each individual barcode reveals that barcodes inserted in early replicating loci tend to be more highly expressed than barcodes inserted in late replicating loci **(Sup. Fig. 2E)**. However, there are still a number of insertions in late replicating loci that appear to be highly expressed. Interestingly, there appears to be no correlation between the levels of transcription originating from a given insertion and changes in the RT of the insertion site **(Sup. Fig. 2E)**. We conclude that, with rare exceptions, transcription from the hPGK promoter fails to advance RT at many ectopic sites in mESCs.

### Transcription of a long transcript enhances the ability of promoters to advance RT

A previous study in chicken DT40 cells provided evidence suggesting that longer transcripts are more effective at advancing RT but there was not a direct comparison of transcription and RT at the same genomic location, in the same cell lines (Blin et al., 2019). To directly investigate the influence of gene size on RT, we took advantage of the negative selection marker of our reporter genes (**Fig. 1**), to delete the sequence between each of our inserted promoters and the +1 TSS for the 96kb Ptn gene, thereby bringing the Ptn gene under the transcriptional control of each promoter (**Fig. 3A**). Under these conditions, the TRE promoter (in the presence of Dox) and the hPGK promoter gained the ability to advance RT (**Fig. 3B**). Additionally, the CAG promoter led to a more substantial advance in RT, whereas the mPGK promoter advanced RT to a similar extent driving either a small or large gene (**Fig. 3B**). Quantification of the levels of Ptn transcription and the RT advance of the Ptn domain induced by each promoter reveals a correlation between RT advance and transcription rates (**Fig. 3C,D, Sup. Fig. 3**). Thus, our results support a role for the length and strength of an active transcription unit in its ability to advance RT, although they still cannot rule out effects due to the nature of the promoters themselves.

**Figure 3:**
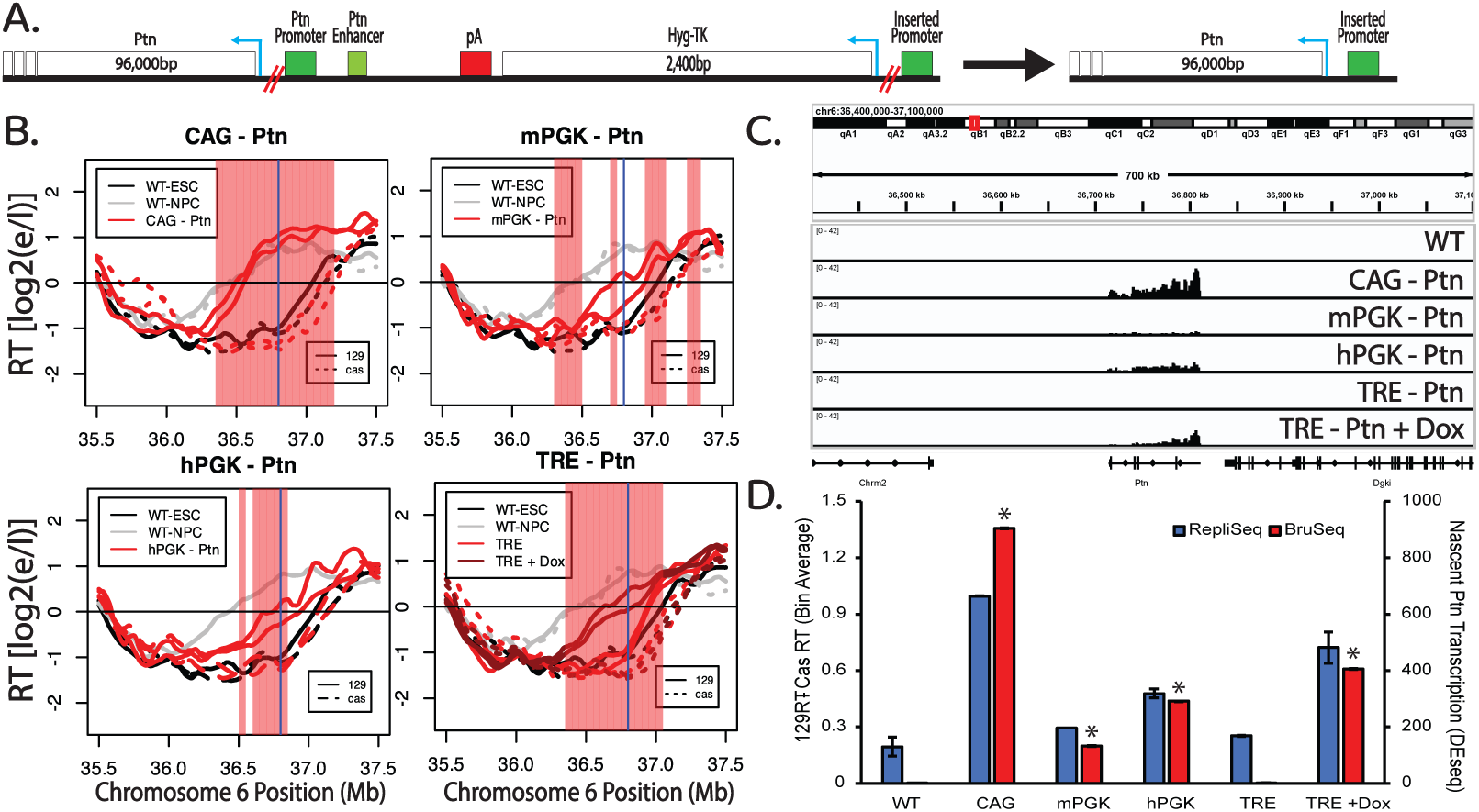
Transcription of a long transcript enhances the ability of promoters to advance RT. A) Schematic of the region directly upstream of the Ptn gene. Deletions of the HTK selectable markers and the endogenous promoter and enhancers of the Ptn gene were made using CRISPR-cas9 targeting the loci marked by the double red lines. Blue arrows indicate the direction of the HTK and Ptn genes. B) RT of the Ptn domain for the CAG-Ptn (top left), mPGK-Ptn (top right), TRE-Ptn (bottom left) and hPGK-Ptn (bottom right) cell lines, presented as in Fig. 1C. C) Genome browser view of nascent RNA signal of the Ptn gene for each one of the four promoters. Replicate experiments are shown in Supplemental Figure 3. D) Bar graph of the relationship between nascent Ptn transcription and the observed advance in the RT of the Ptn domain. The asterisks indicate a significant difference in the levels of Ptn expression calculated using DEseq, relative to Ptn expression from WT 129 allele, which is expected to be zero. The RT advance of the Ptn domain is measured as the average difference between the RT of 50kb bins of the *musculus* allele and the *castaneus* allele. As indicated, the *musculus* (129) allele replicates slightly earlier than the *castaneus* allele in WT; RT differences between the two alleles are normalized for these allelic differences. The error bars show the range of the difference among the 50kb bins between the two replicate experiments.

### Dox-Induced RT advance is rapid, reversible and correlates to transcription levels

The ability of the TRE to advance RT in a Dox-dependent manner when driving the expression of the 96kb Ptn gene provided the opportunity to directly investigate the effect of transcription rate on RT, modulating only a single variable at a time: either the dosage or the time of rtTA DNA binding activity induced by Dox. We treated TRE-Ptn cells with various concentrations of Dox for 24 hours (approximately 1.5 cell cycles), followed by Repli-Seq and Bru-Seq. Under these conditions, increasing concentrations of Dox lead to increased rates of Ptn transcription correlating with advances in RT, until the point at which both plateau (**Fig. 4A-C, Sup. Fig. 4**).

**Figure 4.**
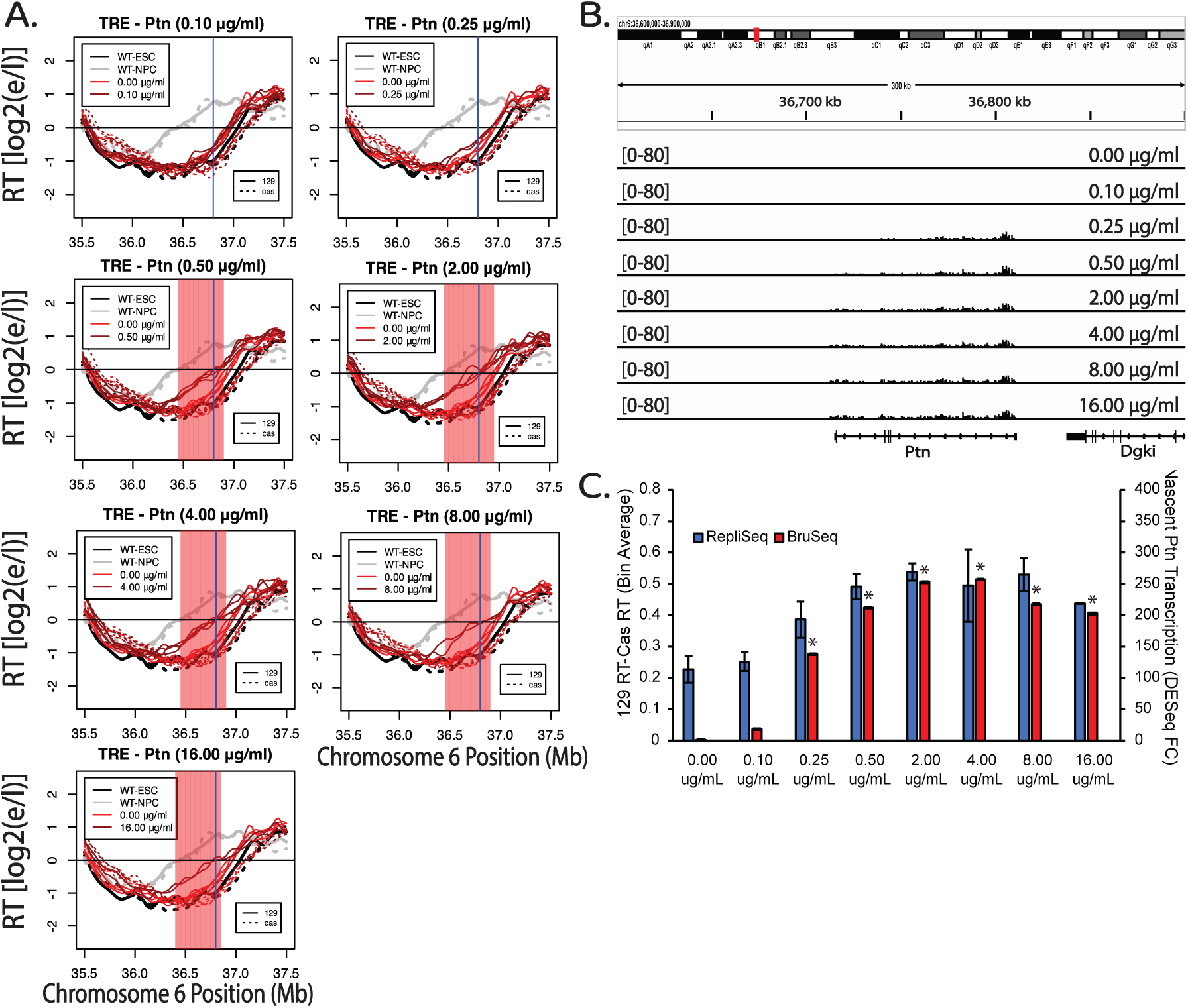
Dosage-dependent advance of RT in response to transcriptional induction of the Ptn gene. A) RT plots of the Ptn domain when TRE-Ptn cells are treated with increasing concentrations of Dox for 24 hours, presented as in Fig. 1C. B) Genome browser view of nascent RNA signal of the Ptn gene at each Dox concentration. Replicate experiments are shown in Supplemental Figure 4. C) Bar graph of the relationship between nascent Ptn transcription and the observed advance in the RT of the Ptn domain. The asterisks indicate a significant difference in the levels of Ptn expression calculated using DEseq, relative to Ptn in a WT 129 allele. The RT advance of the Ptn domain is measured as the average difference between the RT of 50kb bins of the *musculus* allele and the *castaneus* allele. As indicated, the *musculus* (129) allele replicates slightly earlier than the *castaneus* allele in WT; RT differences between the two alleles are normalized for these allelic differences. The error bars show the range of the difference among the 50kb bins between the two replicate experiments.

To measure kinetics of the transcription-induced RT switch, we treated TRE-Ptn cells with 2µg/ml of Dox for 0, 3, 6, 12 and 24 hours post treatment, followed by a 2-hour BrdU (Repli-Seq) or 30-minute BrU (Bru-Seq) label in the continued presence of Dox. Interestingly, 3 hours of Dox treatment was sufficient to induce maximal Ptn transcription rates, and led to a substantial advance in RT, while 6 hours of treatment was sufficient to induce an RT advance similar to 12 and 24 hours of treatment (**Fig. 5A-B**). Taking into account the 2-hour BrdU label following the 6-hour induction, that S phase is approximately 8 hours, and that the average doubling time is approximately 14 hours, we conclude that the RT shift occurs within the first cell cycle of transcriptional induction. We further investigated whether the observed RT advance is sustained upon Dox removal, perhaps as an autonomous epigenetic memory. We treated TRE-Ptn cells with 2ug/ml of Dox for 24h and then removed the Dox for a further 24 hours. Within 24 hours of the removal of Dox, transcription of the Ptn gene was eliminated and the RT of the Ptn locus returned to its original timing (**Fig. 5C-D, Sup. Fig. 5**). Quantification of the levels of Ptn expression and changes in RT show a positive correlation of the increase in Ptn transcription with RT advance during the time course, and a complete elimination of both the Ptn transcription and the RT advance upon removal of Dox (**Fig. 5E, Sup. Fig. 5**). We conclude that the TRE promoter-induced RT shift is rapid and reversible.

**Figure 5.**
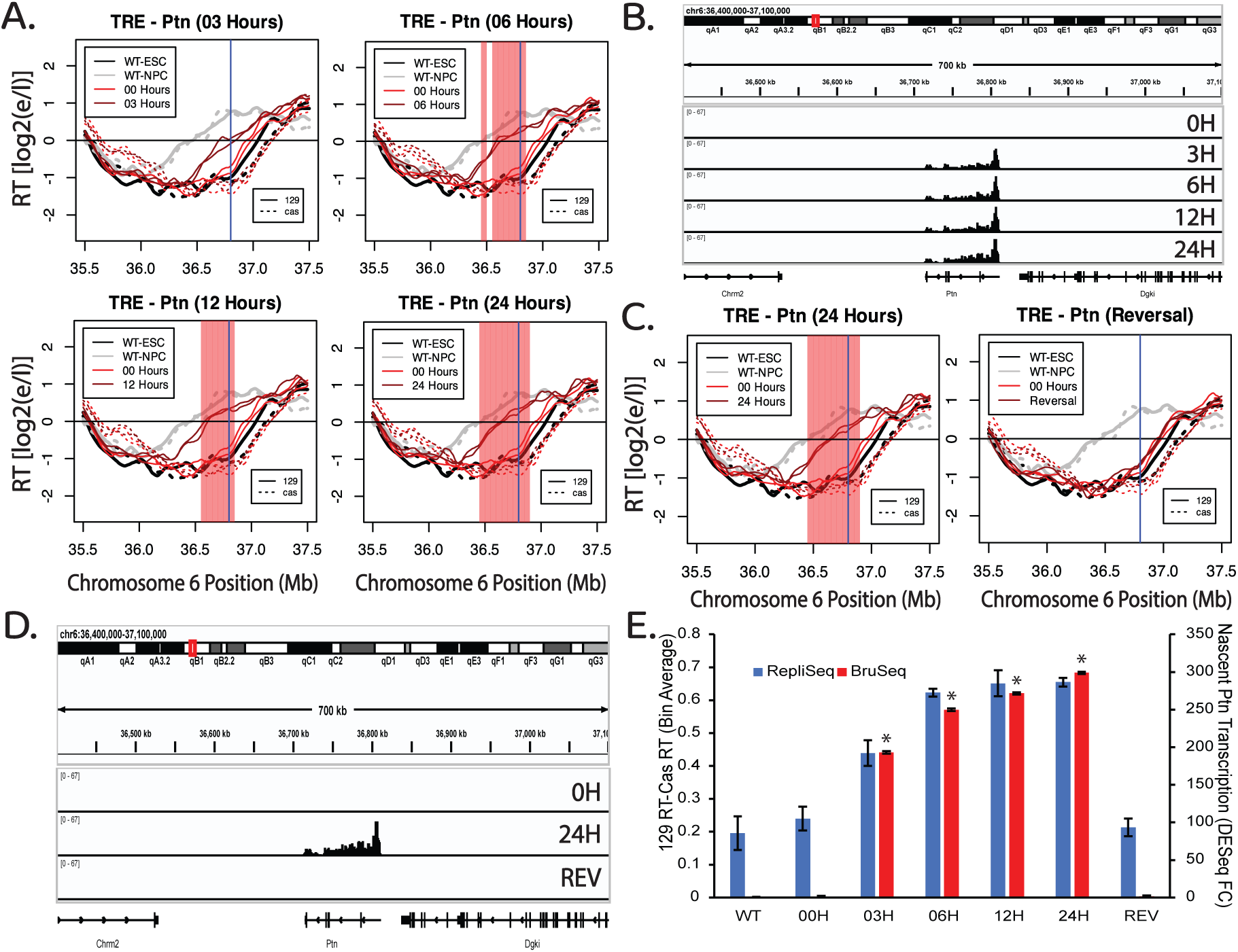
The RT advance induced the TRE-mediated transcription of the Ptn gene is rapid and reversible. A) RT plots of the Ptn domain at different time points after the addition of Dox. BrdU (2 hours) or BrU (30 minutes) was added at the end of each Dox incubation time period in the continued presence of Dox. In each plot, the black and gray lines indicate the RT of WT ESCs and NPCs respectively. The red lines indicate the RT of the insert cell line with (dark red) or without (light red) Dox. Solid and dashed lines indicate the RT of the *musculus* and *castaneus* alleles respectively. Vertical blue line indicates the +1 of the Ptn gene. The red shading indicates 50kb windows with statistically significant differences in RT between WT 129 and the modified 129. B) Genome browser view of the Ptn domain displaying nascent RNA signal of the Ptn gene at each time point. Replicate experiments are shown in Supplemental Figure 5. C) RT plots of the Ptn domain after 24 hours of Dox treatment and subsequent removal of Dox for an additional 24 hours. In each plot, the black and gray lines indicate the RT of WT ESCs and NPCs respectively. The red lines indicate the RT of the insert cell line with (dark red) or without (light red) Dox. Solid and dashed lines indicate the RT of the *musculus* and *castaneus* alleles respectively. Vertical blue line indicates the +1 of the Ptn gene. The red shading indicates 50kb windows with statistically significant differences in RT between WT 129 and the modified 129. D) Genome browser view of the Ptn domain displaying nascent RNA signal of the Ptn gene at each condition in Fig. 5C. E) Bar graph of the relationship between nascent Ptn transcription and the observed advance in the RT of the Ptn domain. The asterisks indicate a significant difference in the levels of Ptn expression calculated using DEseq, relative to Ptn in a WT 129 allele. The RT advance of the Ptn domain is measured as the average difference between the RT of 50kb bins of the *musculus* allele and the *castaneus* allele. As indicated, the *musculus* (129) allele replicates slightly earlier than the *castaneus* allele in WT; RT differences between the two alleles are normalized for these allelic differences. The error bars show the range of the difference among the 50kb bins between the two replicate experiments.

### Transcription of the Ptn Gene is not Necessary to Advance RT During Differentiation

The results we have shown with TRE-driven short and long transcripts demonstrate directly that, when isolated as the only variable, transcription of the Ptn gene can be sufficient to advance RT. The fact that a small TRE reporter gene cannot advance RT with the same promoter at the same location eliminates the possibility that binding of the rtTA transactivator on its own is sufficient to advance RT but, rather, the effect on RT requires a sufficient length and strength of transcription. This raises the question as to whether the natural RT advance that accompanies Ptn induction during differentiation requires Ptn transcription. We identified the Ptn domain as an RT-switching domain over 20 years ago (Hiratani, PNAS 2004), and showed that its RT is advanced coincident with movement away from the nuclear periphery and activation of transcription (Hiratani et al., 2010). The coincidence was such that we were not able to delineate the order in which these correlated events occurred over the course of *in vitro* differentiation, consistent with a direct link between transcription and RT. To directly address the causality between transcription and RT during differentiation, we deleted both the promoter and predicted enhancers for Ptn transcription, either on the *musculus* (129) allele alone or on both *musculus* and *castaneus* alleles, and differentiated these cells to neural precursor cells (NPCs). Results (**Fig. 6**) demonstrated that the RT advance during differentiation still occurred despite elimination of detectable Ptn transcription or whether detectable transcription was eliminated on one allele or both alleles. In the heterozygote configuration, both alleles were found to advance similarly, despite the differential presence of Ptn gene induction. Thus, Ptn transcription is sufficient to advance RT in mESCs, but not necessary to advance RT during differentiation. Given that Ptn was the only RNA detected throughout this domain during NPC differentiation (**Fig. 6C**) by total, rRNA-depleted RNA-seq, and has previously been shown to be the only transcriptional activity detected throughout this domain in either mESCs or mNPCs (Giorgetti et al., 2016; Rivera-Mulia et al., 2018), there is no evidence for a requirement of any transcription for an RT shift during differentiation within this developmentally RT-switching RD.

**Figure 6.**
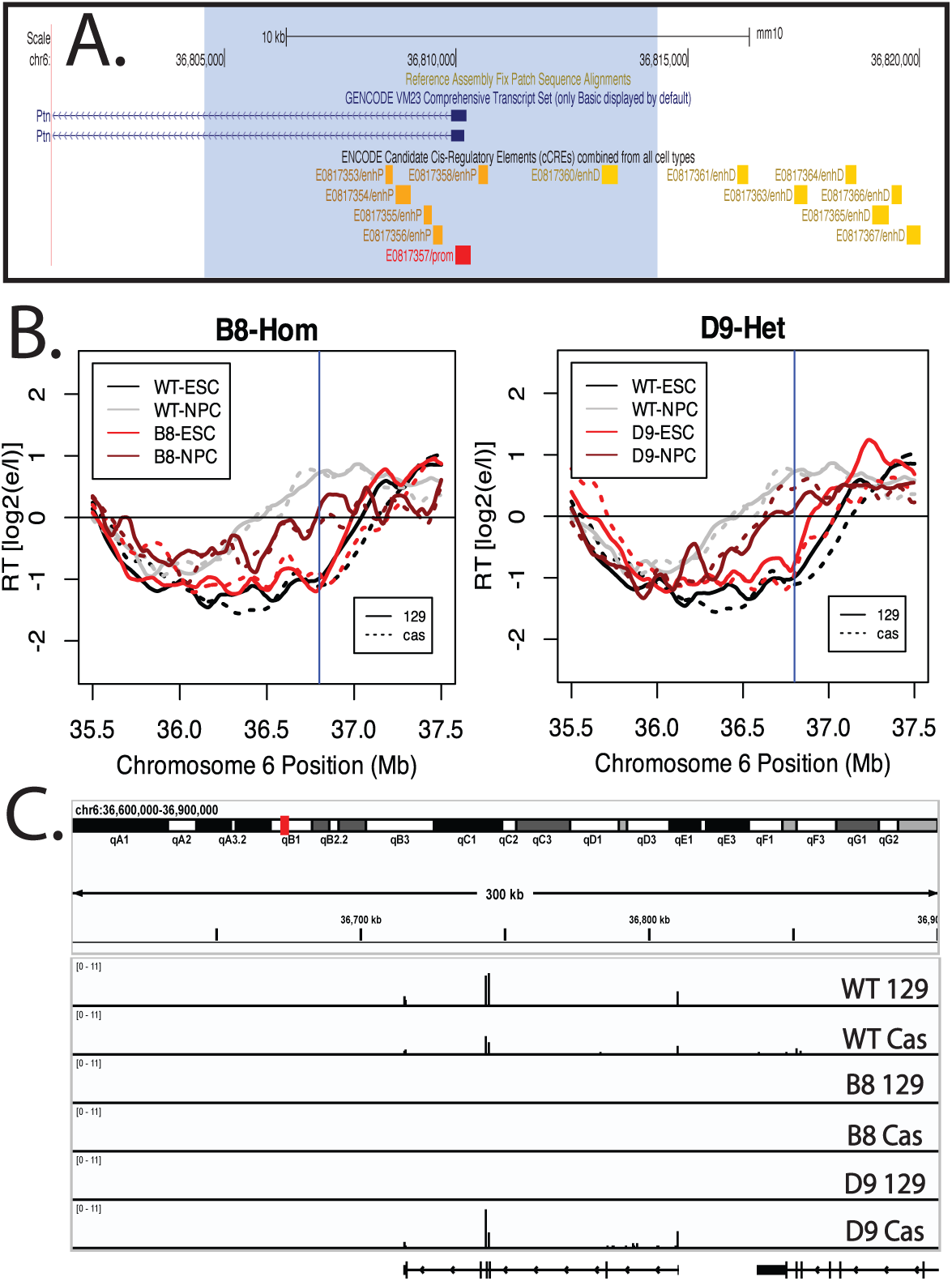
Deletion of the Ptn promoter and elimination of Ptn transcription does not significantly affect the RT advance of the Ptn domain during NPC differentiation. A) UCSD Genome Browser track indicating the part of the Ptn domain that was deleted. The deleted sequence is highlighted by the blue shade. Candidate promoter, proximal enhancers and distal enhancers are indicated by the red, orange and yellow boxes. B) RT plots of clones B8 and D9, in which the Ptn promoter deletion is homozygous and Heterozygous respectively. Dark gray and light gray indicate the RT profiles of WT ESCs and NPCs respectively. Light red and dark red indicate the RT profiles of ESC and NPC deletion clones respectively. Solid and dashed lines indicate the RT of the musculus and castaneous alleles respectively. Vertical line indicates the site of the deletion. C) Browser track of total rRNA-depleted RNA-seq for WT NPCs and the two deletion clones. RNA-seq was performed in lieu of Bru-Seq because cell death during the differentiation process makes it difficult to collect enough cells to perform Bru-Seq.

## DISCUSSION

The RT program is essential for maintenance of the global epigenome, strongly correlated with other structural and functional features of chromosomes and is regulated during development. Therefore, it is likely that changes in RT during differentiation are an integral part of organismal development, but we have little understanding of how they are executed. One longstanding hypothesis has been that transcription is sufficient for early replication and potentially drives RT changes during development (Goldman et al., 1984; Hatton et al., 1988; Therizols et al., 2014). If true, then such changes, affecting large replication domains and in some cases multiple genes, could alter domain-wide epigenome architecture to assemble a chromatin environment more robust for transcriptional throughout the domain (Klein et al., 2021). As attractive as this model is, it is challenged by both manipulative studies and genomics data that provide numerous contradictory examples of late replicating and expressed genes discussed in the introduction (Blin et al., 2019; Hassan-Zadeh et al., 2012). Here, we have taken a reductionist approach to systematically alter single variables that could reconcile these contradictions. We find that transcription can be sufficient to advance RT depending upon the length and strength of the resulting transcript; small genes can be transcribed with no detectable effect on RT but when driven by promoters that drive high levels of transcription, can advance RT. Large transcription units can advance RT even at low levels of transcription and show an RT shift that is positively correlated with transcription rate. However, we also show that transcription is entirely dispensable for developmental RT shifts, demonstrating that non-transcriptional mechanisms can be equally effective at driving an RT advance. Together, these experimental results provide an empirical rationale to reconcile data in the literature relating transcription and RT.

### Length and Strength Explain Much of the Literature with Ectopic Transcription

Our results provide a solid empirical base with which to re-evaluate many decades of literature by considering length and strength of transcription as sufficient but not necessary to advance RT. This seems particularly true for the ectopic vector induction literature (see Introduction). Small vectors may or may not advance RT, while long transcripts are more effective at doing so and ectopic inserts that do not induce transcription can advance RT through alternative mechanisms. Our results extend very interesting observations in chicken DT40 cells (Blin et al., 2019). While this study was focused on the effects of RT on fragile site expression, and the design of the experiments did not directly compare transcription length, strength and RT at the same locus, their results were consistent with those reported here. An apparent contradiction with our results was that Blin et. al. did not find any significant effect of transcriptional induction by the TRE promoter even when driving a long transcript. There are several differences in our approaches that could account for this discrepancy. First, Blin et. al. detected a small advance in RT prior to induction and did not titrate their induction to determine whether they had achieved maximum transcription thereafter, thus it is possible that they had leaky transcription prior to induction and/or did not fully induce the TRE promoter. Second, steady state transcription was measured rather than nascent transcription rates as we have done. Third, RT relative to the wild-type locus was measured in different cell lines; our direct allelic comparison in a hybrid-genome cell line offers much improved sensitivity to detect RT changes. Perhaps most importantly however, RT and transcription were assayed by single locus PCR quantification, which in our hands can be misleading. Here, we have performed genome-wide analyses to capture whole-domain RT and all nascent transcription, with and without induction, at different times and concentrations of inducer, and relative to an internal wild-type control allele. Thus, our results have the potential to explain all of the discrepancies in the literature regarding ectopic manipulations of genomic sites to study the relationship of RT to transcription.

While our results to not unveil detailed mechanisms by which transcription can advance RT, they do focus our attention on the act of transcription of a sufficient length of chromatin. Most compellingly, we show that an artificial trans-activator, as the only variable, can be targeted to a small DNA sequence to induce transcription of a small gene with no effect on RT, while the same targeting event to the same small DNA sequence, when inducing a larger gene, substantially advances RT at the same or even lower targeting dosage. It is therefore unlikely that the effects on RT are indirectly related to Tet trans-activator binding and its downstream effects on local chromatin or its recruitment of the transcription machinery per sé. Rather, the RT advances must be a consequence of transcribing a larger segment of chromatin. Further, the fact that the RT advance can occur within a few hours and is rapidly reversible suggests a direct dependence upon transcription itself. Due to the short duration of the mESC G1 phase, we unfortunately cannot conclude whether or not transcriptional induction must occur prior to mitosis or the early G1 phase Timing Decision Point (Dimitrova & Gilbert, 1999). This will require recapitulating the system in cell lines with longer G1 phases. When considering the short transcripts, our data demonstrate that promoters that advanced RT drove detectable, albeit low level, read-through transcription beyond the vector polyA sequence. Thus, it is possible that their ability to advance RT is not directly due to transcription rate but to the frequency of long transcription. Altogether, the effects of ectopic insertions of promoters on RT may only indirectly depend upon the nature of the promoter but, rather, depend solely on the length and strength of transcription.

In contrast to the literature on ectopic manipulation of genomes, what is not fully explained are genomics results surveying natural relationships between RT and transcription in different tissues and differentiation systems (see introduction). There are still numerous examples of large to very large, even highly expressed, late replicating and expressed genes and rare examples of induced genes experiencing delays in RT and genes that are only transcribed when late replicating (Heskett et al., 2022; Rivera-Mulia et al., 2015). Many of these datasets score only steady state mRNA or steady state total RNA and most analyses do not take into account the effects of neighboring genes within the same domains. What is needed is a thorough re-analysis of genome-wide RT and nascent total transcription datasets to test the length and strength hypothesis. It also remains possible that the rules of engagement are different for ectopic vs. natural transcriptional induction.

### A system to rapidly and reversibly alter RT opens up avenues to explore mechanism

Our finding of a system for inducible RT shifts opens up a host of experimental approaches to investigate mechanisms linking transcriptional induction to RT changes. How transcription mediates this change remains a subject of speculation. One possibility is that transcription of large genomic segments could move pre-loaded, and remarkably stable Mcm helicases (Kuipers et al., 2011), in such a way that concentrates them downstream of the transcription unit, sufficiently to create a high efficiency initiation site (Sasaki et al., 2006). What are needed to test this hypothesis are effective means to map the positions of Mcm double hexamers in living cells (Li et al., 2023). Another possibility is that longer transcripts tend to be spliced, and it is well established that proximity to speckles, the sub-nuclear storage compartments for splicing machinery is highly correlated with early replication and splicing may induce movement of domains towards the speckles (Bhat et al., 2024; Chen et al., 2018). Our results underscore the need for a focus on transcript length and strength as a mechanism driving RT changes.

### Transcription is not necessary to advance RT during a cell fate change

With respect to natural RT shifts in the context of differentiation, we demonstrate that transcription is not necessary for a differentiation-induced RT advance. Some other mechanism, perhaps developmentally-activated ERCEs present in the Ptn domain (Sima et al., 2019), can elicit the RT switch in the absence of transcription. These two mechanisms could interact to form a positive feedback loop if either transcription or ERCEs could initiate an RT switch, followed by either binding of core TFs to activate latent ERCEs or a higher level of transcriptional activation following an RT shift, respectively (Turner et al., 2024). Together, this could serve to lock in epigenetic states in a step-wise fashion to provide flexible alternatives during a cell fate transition. Experimental approaches that can uncouple transcription induction from core TF binding during a natural cell fate transition will be needed to investigate the separate roles of these different regulatory mechanisms.

### Reporter Genes That do not Affect RT Provide Useful Tools

Our work also identified promoters that can drive ectopic gene expression in either a constitutive or inducible fashion, and function well as selectable markers, without affecting RT. Since RT is so closely related to sub-nuclear position and chromatin compartment, relationships that are also poorly understood, these markers may prove useful in applications where integration is desired without affecting RT, for example, to study large-scale chromosome structure and function (4D Nucleome Consortium et al., 2024; Ryba et al., 2010; Wang et al., 2021).

## MATERIALS AND METHODS

### Cell Culture

F121:R26puroR-M2rtTA mESCs (Quinodoz et al., 2021), expressing the tetracycline transactivator from the Rosa22 locus, and all mutant cell lines derived from the parental cell line, were cultured, maintained, and passaged according to the 4DNucleome SOP (https://data.4dnucleome.org/protocols/cb03c0c6-4ba6-4bbe-9210-c430ee4fdb2c/) for F121:R26puroR-M2rtTA unless otherwise stated. For Dox-induction, doses ranged from 0 to 16ug/ml of Dox for 24 hours. Higher doses of Dox led to significant rates of cell death making it challenging to collect enough material for RT and transcription assays. ESCs were differentiated into NPCs using the 4DNucleome differentiation protocol (https://data.4dnucleome.org/protocols/fa439403-03b2-4ef4-be5f-3c76fb2e8e1a/).

### Ectopic Insertions

F121:R26puroR-M2rtTA mESCs were nucleofected with a donor plasmid containing one of the four promoters used, a hygromycin resistance/thymidine kinase fusion gene and a transcription termination sequence, flanked by homology arms targeting sequences directly upstream of the Ptn promoter on chromosome 6. The donor plasmid was co-transfected with two modified Px458 cas9 plasmids containing two guide RNAs targeting sequences upstream of the Ptn promoter, and a plasmid expressing mCherry used to assess the success of the nucleofection. Nucleofected cells were cultured for two days and then treated with 350ug/mL of Hygromycin. Colonies of hygromycin resistant clones were isolated and plated into 96-well plates 7-10 days after the addition of Hygromycin. Colonies were expanded into multiple 96-well plates and screened by PCR with a set of primers, one targeting the genomic sequence flanking the homology arms and the other targeting the inserted sequence. A different set of screening primers was used for the insertion of the CAG promoter due to difficulties with the PCR amplification of the CAG promoter. Clones were also screened with a wild type set of primers, one targeting the genomic sequence flanking the homology arms and the other targeting the sequence between the two guide RNAs. A positive on both screens indicated that the insertion was made on a single copy of chromosome 6.

### Deletions

Cell lines with ectopic insertions were nucleofected with two modified Px458 cas9 plasmids containing two guide RNAs, one targeting a sequence between the inserted promoter and the inserted hygromycin resistance gene and the other targeting a sequence directly upstream of the +1 of the Ptn gene. Two days after the nucleofection, cells were treated with 100μg/ml of Ganciclovir. Colonies of Ganciclovir resistant clones were isolated and transferred to 96 well plates 10-14 days post treatment. Clones were expanded into multiple 96 well plates and screened with a set of primers, one targeting the inserted sequence upstream of the guide RNA targeted sequence and the other targeting a genomic sequence downstream of the +1 of the Ptn gene.

### E/L Repli-Seq

E/L Repli-Seq was performed as previously described (Marchal et al., 2018). Briefly, cells were labeled with BrdU for 2 hours and then fixed in 75% cold ethanol. Fixed cells were FACS sorted into early-S and late-S fractions based on DNA content. Genomic DNA, from 40,000 cells per fraction, was isolated, sheared, and adaptor-ligated with NEBNext Ultra Library DNA Prep Kit. These adaptor-ligated DNA fragments were enriched for nascent DNA via BrdU immunoprecipitation, PCR amplified, indexed, and sequenced on HiSeq2500 or NovaSeq6000 aiming to obtain ≥30 million clusters of ≥50 bp reads per library.

### E/L Repli-Seq Analysis

Raw sequencing reads were aligned to an mm10 reference genome with bowtie2 and subsequently musculus and castaneous specific reads were isolated using SNPsplit (Krueger & Andrews, 2016). Aligned and parsed reads were then quality filtered and depleted of PCR duplicate reads. To calculate the statistical significance of RT differences across samples, processed early and late fraction reads were binned intp 50kb windows, genome-wide and the log2 ratio of early to late fraction was calculated for each window, which was then quantile normalized to the RT of WT F121 cells. Statistical significance of RT changes for all windows in each sample, relative to WT, were calculated using RepliPrint (Ryba et al., 2011). To generate RT plots, reads were binned into 5kb windows, and the early-to-late log2 ratio was quantile normalized to the RT of WT F121 cells, scaled and smoothed in 300kb bins. It is important to note that scaling, which was applied on the data to generate RT plots but not for calculating statistical significance, alters the dynamic range of the data (∼-3 - ∼3) from the original dynamic range (∼-7 – 7). This means that RT values in RT line plots are not comparable to the RT values in the rest of the plots.

### Bru-Seq

Bru-Seq libraries were prepared according to previously published protocols (Paulsen et al., 2014).Cells were labeled for 30 min with Bromouridine (Sigma-Aldrich, 850187). BrU labeled cells were collected and total RNA was isolated from the cells using the Direct-zol RNA Miniprep Plus kit (Zymo, R2070). BrU labeled RNA was isolated using a Bromodeoxyuridine antibody (Sigma-Aldrich, B5002) conjugated to goat anti-mouse IgG Dynabeads (Thermo-Fisher, 11033). Nascent BrU-labeled RNA was fragmented at 85°C for 10min and reverse transcribed into cDNA using NEBNext Ultra II RNA First Strand Synthesis Module (NEB, E7771) and the NEBNext Ultra II Directional RNA Second Strand Synthesis Module (NEB, E7550). Libraries were prepared from cDNA using the NEBNext Ultra II Library Prep kit (NEB, E7645), indexed and sequenced on NovaSeq6000 to obtain ≥80 million of 100 bp single end reads.

### Bru-Seq Analysis

Raw sequencing reads were aligned to the mouse ribosomal DNA with bowtie2, keeping only reads that did not align to the ribosomal DNA. Non-ribosomal reads were subsequently sequenced to the mm10 genome, using tophat aligner and the following flags {--min-isoform-fraction 0 --max-multihits 1 --no-coverage-search --library-type fr-unstranded --bowtie-n --segment-mismatches 1 --segment-length 25}. The Bru-Seq data for the cell lines with the HTK insertions were aligned to a custom genome in which the Mitochondrial DNA sequence was replaced with the HTK sequence. Allele specific reads were isolated using SNPsplit. Bru-Seq tracks for visualization were generated using the STAR RNA-seq aligner (Dobin et al., 2013). Counts tables were generated using featureCounts (Liao et al., 2014). Differential expression analysis was conducted using the R package DESeq2.

### RNA-seq

From 300ng total RNA extracted using Direct-zol RNA MiniPrep (Zymo research R2050), ribosomal RNA was depleted using NEBNext® rRNA Depletion Kit v1 (NEB E6310). The entire resulting RNA was used for 1st strand synthesis by NEBNext Ultra II RNA First Strand Synthesis Module (NEB, E7771), following “Directional RNA-seq Workflows”. Second strand was synthesized using NEBNext Ultra II Directional Second Strand Synthesis Kit (NEB E7550) then cDNA was made into sequencing library using NEBNext Ultra II DNA Library Prep Kit for Illumina (NEB E7645).

### RNA-seq Analysis of NPC Data

RNA-seq reads were aligned to an mm10 genome with masked 129/Cas SNPs, using the STAR RNA-seq aligner (Dobin et al., 2013). Reads aligned to mm10 were subsequently parsed into 129 and Cas specific reads using SNPsplit. Browser tracks were generated using the inputAlignmentsFromBAM function of the STAR RNA-seq aligner (Dobin et al., 2013).

#### CM Insertion RNA-seq Data Analysis

Barcodes were matched to 61/107 insertions in the CM1417 and CM1420 clones. To assess the abundance of these barcodes in the RNA-seq datasets, reads containing the exact 16 nucleotide barcode were extracted from the raw FastQ files and counted. Barcode counts were normalized by the total number of sequencing reads of each replicate (RPM). Separately, to assess the average levels of GFP expression per insertion in each clone, reads were mapped onto a custom reference genome containing the sequence of the GFP gene in chromosome M, using the STAR RNA-seq Aligner (Dobin et al., 2013). Count tables for each clone were prepared using featureCounts (Liao et al., 2014). The RPKM of the average counts across two replicates was calculated using the EdgeR R package (Robinson et al., 2010). The RPKM of GFP for each clone was divided by the number of insertions present in that clone. Subsequently, the log2 of the RPKM for each gene was calculated.

## ACKNOWLEDGEMENTS

We thank B. Washburn and C. Pye for their valuable input in the design of the constructs and the guides used in these experiments. We also thank C. Vied for her valuable input with sequencing our libraries. We thank A. Chow and M. Guttman for sharing cell line F121:R26puroR-M2rtTA. Finally, we thank B. Chadwick for the modified px458 plasmid used to clone CRISPr Cas9 guide RNA sequences. This work was funded by NIH grant GM083337 to DMG and NIH Common Fund “4D Nucleome” Program grant U54DK107965 (BvS and DMG). The Oncode Institute is partly supported by KWF Dutch Cancer Society.

---

**Supplemental Figure 1:**
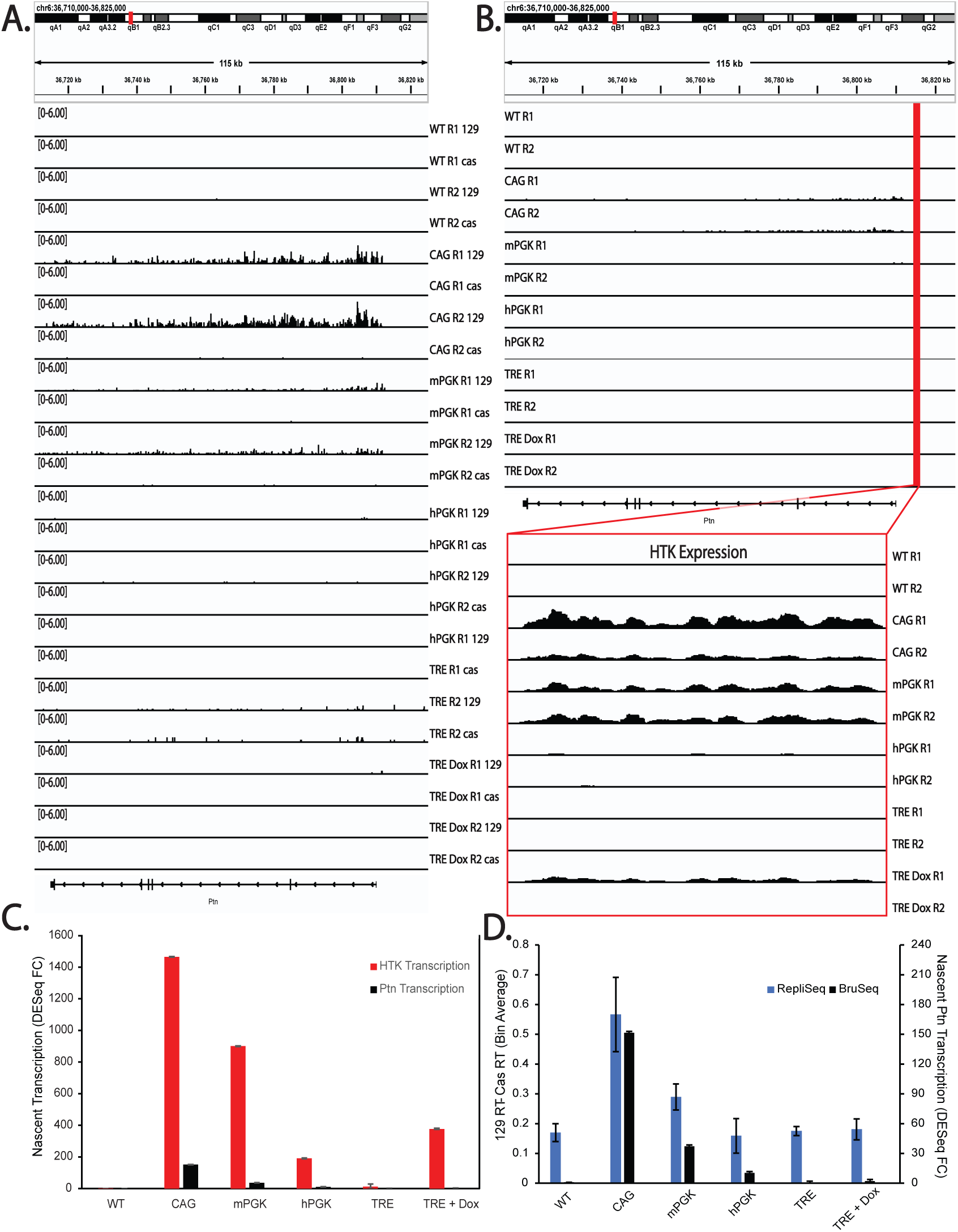
A) Genome Browser tracks of genome-parsed Bru-Seq replicates for the cell lines with the Promoter-HTK insertions (vector sequences not included), indicating that Ptn read through reads originate exclusively from the 129 allele. B) Genome Browser tracks of unparsed Bru-Seq replicates for the cell lines with the Promoter-HTK insertions, showing only the inserted allele for each replicate. Top tracks display the read through expression of the Ptn gene, while the zoomed in, bottom tracks display the expression of the HTK gene. C) Bar graph comparing HTK and Ptn read through expression in the Promoter-HTK cell lines. D) Bar graph of the levels of read through Ptn expression in relation to changes in the RT of the Ptn domain.

**Supplemental Figure 2:**
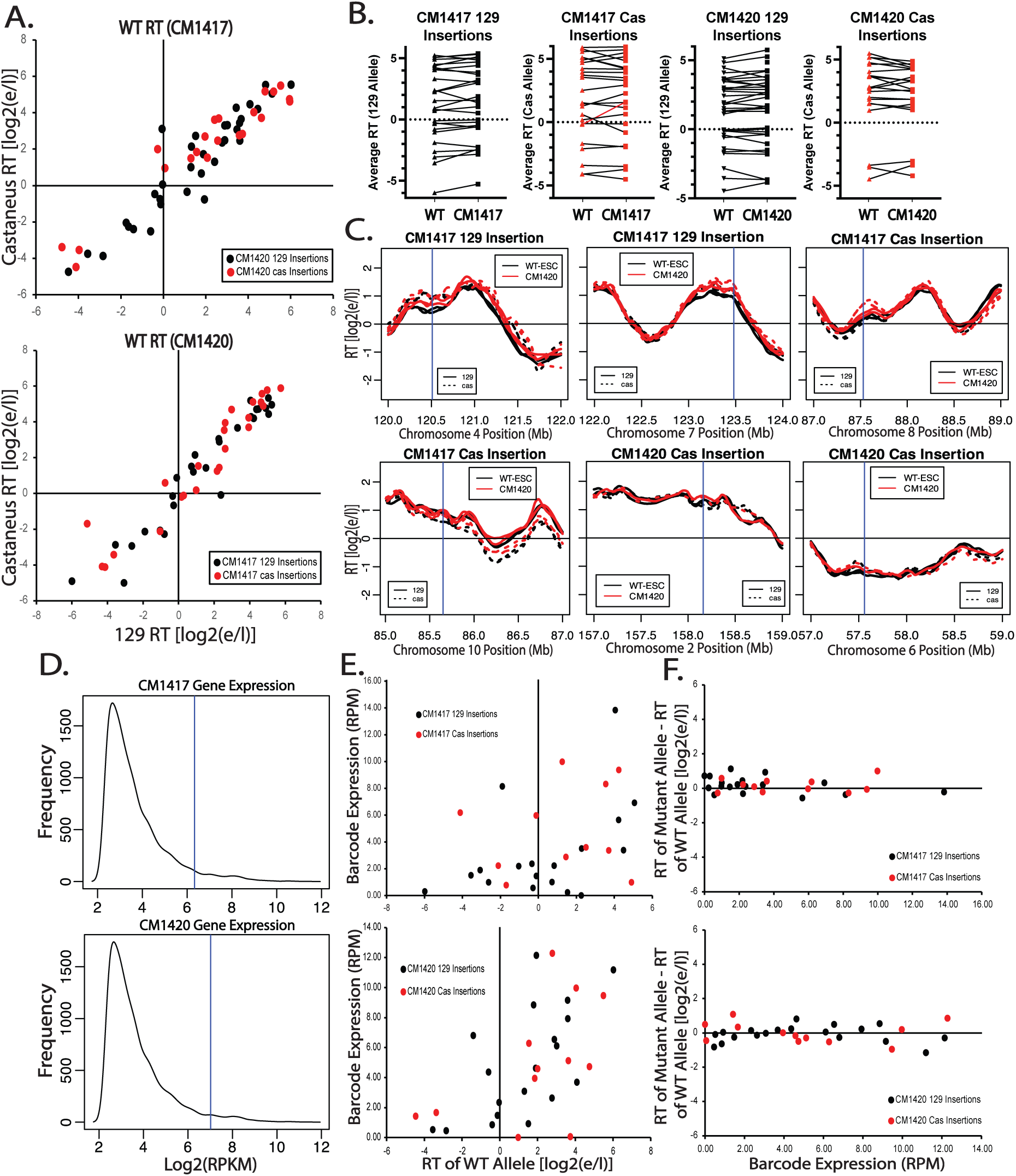
**A)** Scatter plots of the RT (129 vs Cas) of the WT parental alleles of all CM1417 (left) and CM1420 (right) ectopic insertion sites. Black and red dots represent loci where the eventual insertion was made in the 129 and Cas allele respectively. B) RT of the CM1417 129 insertions (left), CM1417 Cas insertions (middle-left), CM1420 129 Insertions (middle-right) and CM1420 Cas Insertions (right) before (WT, triangles) and after (CM1417 or CM1420, squares) the PB insertion. There were no statistically significant advances, calculated using RepliPrint, in RT across both clones, with the exception of the site inserted at Chromosome 8, 87.5Mb, of the Cas allele of CM1417. The locus with the statistically significant RT change is marked with a red line in the CM1417 plot. C) RT plots of six insertion sites where the largest advances in RT of the insertion allele were observed, including the statistically significant RT change at the Cas Chromosome 8 allele (top right). D) Distribution of all expressed genes in CM1417 (top) and CM1420 (bottom) clones. The x-axis represents the log2(RPKM) expression level of each gene and the y-axis represents the number of genes that are expressed at each level. RPKM was calculated from the average read count for each gene across two replicates. Genes with extremely low or no expression (RPKM<5) were excluded from the plot. Average GFP expression, calculated by dividing the GFP RPKM of each clone by the number of insertions in that clone, is marked with vertical blue lines. E) Position effect of each insertion locus on the expression level of each insertion. To differentiate between expression levels at each insert, only reads containing the 16-nucleotide barcode were extracted from the unprocessed FastQ and were used to calculate the RPM for each barcode. F) Effect of transcription of individual insertions on the RT of the insertion site. The Y-axis range represents the dynamic range of RT in each clone.

**Supplemental Figure 3.**
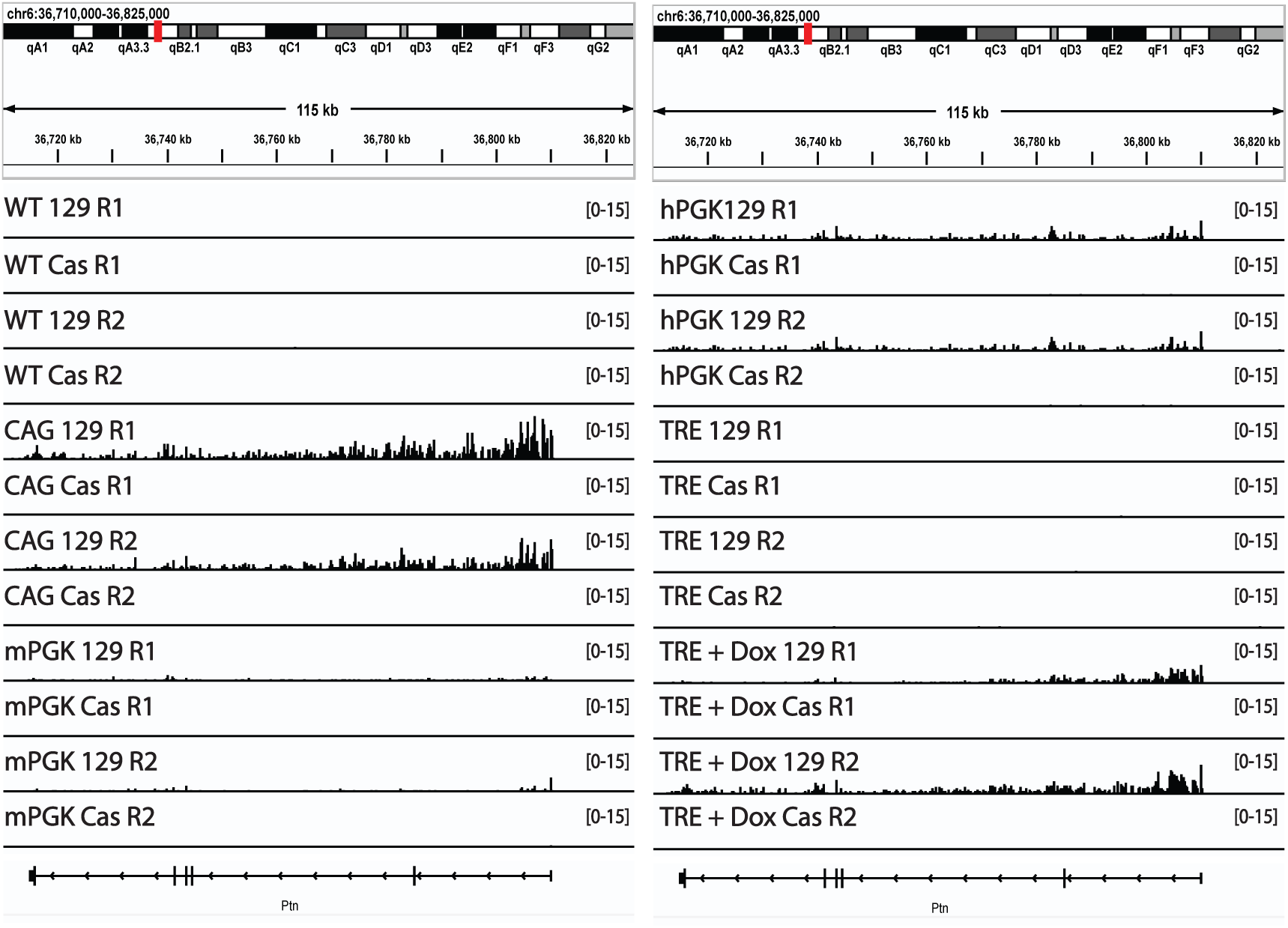
Genome Browser track of parsed Bru-Seq replicates for the cell lines in which the inserted promoters drive Ptn transcription.

**Supplemental Figure 4.**
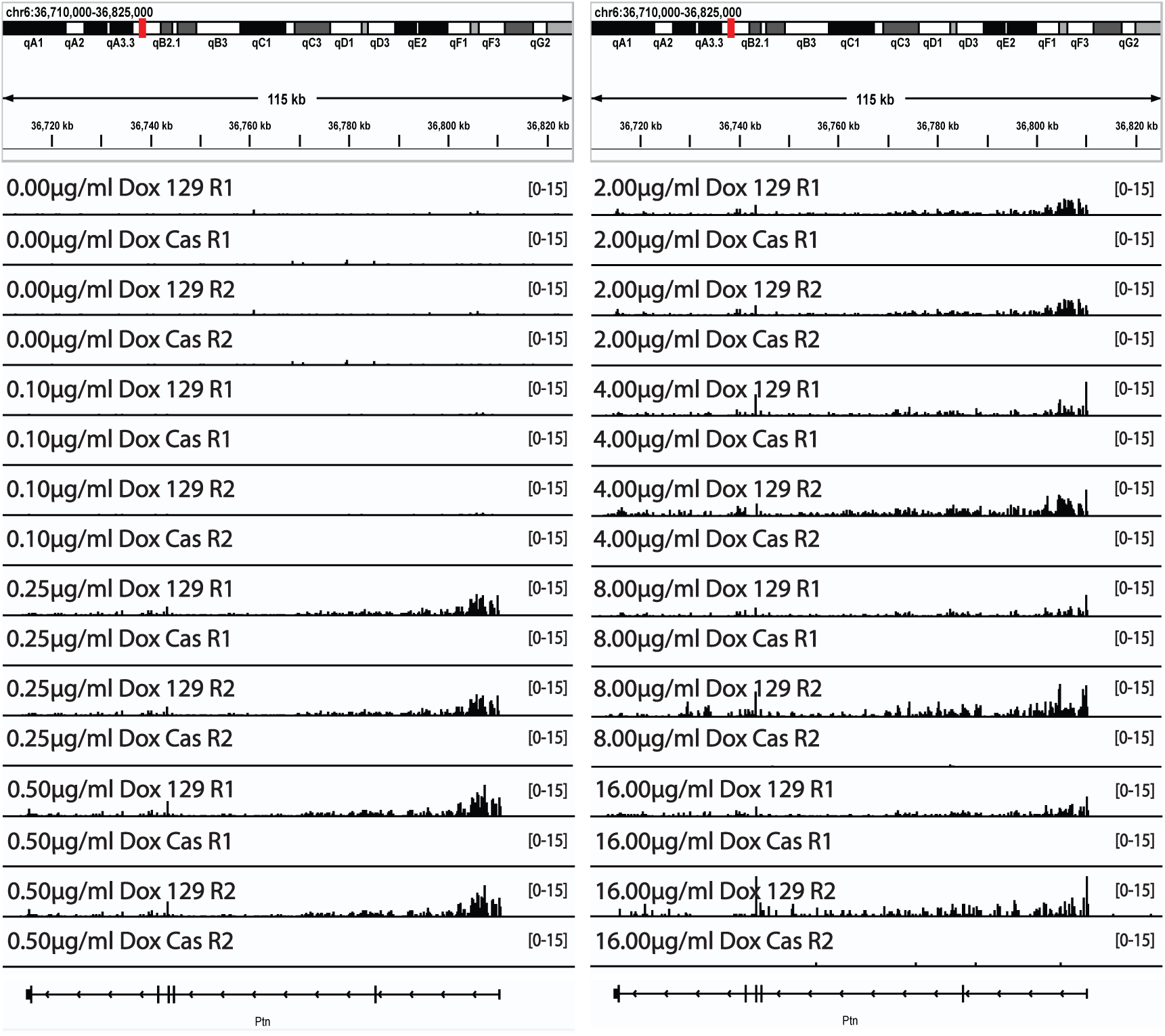
Genome Browser track of parsed Bru-Seq replicates displaying Ptn expression at different concentrations of Dox in the TRE-Ptn cell lines.

**Supplemental Figure 5.**
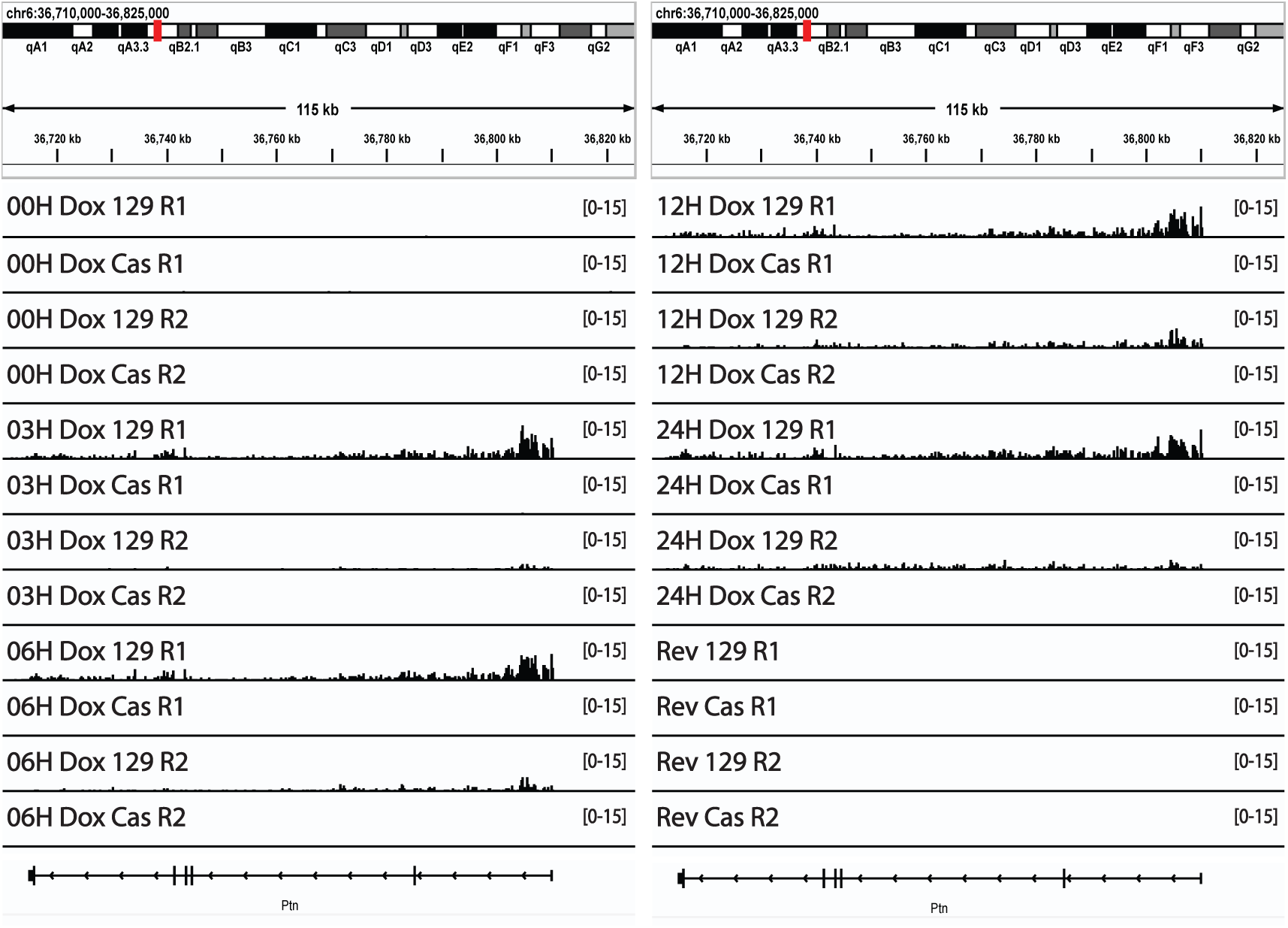
Genome Browser track of parsed Bru-Seq replicates displaying Ptn expression at different time points after the addition of Dox in the TRE-Ptn cell lines.

